# Adapting a commercial anti-*Aspergillus* IgG ELISA kit for penguin sera: A novel approach using anti-chicken IgY antibody for the detection of anti-*Aspergillus* IgY titer

**DOI:** 10.1101/2025.09.08.673393

**Authors:** Takahito Toyotome, Kana Hatazawa, Naoya Matsumoto, Kyogo Hagino, Nanako Sawayama, Nobutaka Sato, Chiharu Tanaka, Megumi Itoh, Kazutaka Yamada

## Abstract

Penguins are susceptible to aspergillosis and are affected by the causative fungal pathogens *Aspergillus* spp., under immunocompromised conditions. Currently, there are limited serological tests available for diagnosing *Aspergillus* infections in penguins, highlighting the need for new diagnostic methods. While an anti-*Aspergillus* IgG detection kit using ELISA is commercially available and widely used in human medicine, it is not applicable for penguins because it incorporates anti-human antibodies as the secondary antibody for detection. To address this issue, an anti-chicken IgY antibody was incorporated into a commercial anti-*Aspergillus* IgG detection ELISA kit. First, anti-chicken IgY antibody was examined for cross-reactivity to penguin IgY, and the antibody recognized IgY from King, Gentoo, and African penguins in sera. Subsequently, serum samples from healthy penguins and penguins with aspergillosis were examined using anti-chicken IgY antibody incorporated into the anti-*Aspergillus* IgG ELISA kit. The results suggest that the combination method is applicable for the detection of anti-*Aspergillus* penguin IgY antibodies. However, because background titers can vary among individuals, we propose that routine monitoring could aid in the early detection of aspergillosis, even before the onset of symptoms.

## INTRODUCTION

*Aspergillus*, a common fungal genus in the environment, is also a pathogenic fungal genus in humans and animals that causes aspergillosis. Aspergillosis has been reported in various avian species; it remains an important concern in the veterinary field [1]. Penguins are extremely susceptible to Aspergillus spp. infection. In particular, under uncomfortable conditions, such as inadequately hot or cold conditions, penguins are even more susceptible to *Aspergillus* [2].

Early diagnosis of aspergillosis in penguins is difficult because of its common symptoms. Our previous research indicated that computed tomography (CT) imaging is useful for diagnosing aspergillosis in penguins [10, 16]. However, CT scanning has some limitations, including specificity for aspergillosis, accessibility to equipment, and high cost. As described in previous publications, the inflammatory response elicited by aspergillosis results in a marked increase in globulin fractions and a decrease in the albumin/globulin (A/G) ratio in sera [7, 12]. However, this elevation is not specific to aspergillosis.

Various serological tests have been developed to diagnose human aspergillosis. Some of them, such as an immunodiffusion test and *Aspergillus* antigen (galactomannan) enzyme-linked immunosorbent assay (ELISA) kit [6, 8], and *Aspergillus* lateral-flow devices [11] are applicable to aspergillosis in animals, including penguins. However, most of them are not available, and their use for aspergillosis in animals is uncertain. Immunodiffusion method is one method that may be useful in diagnosing aspergillosis in penguins [4]. Immunodiffusion testing has replaced worldwide with kits that determine anti-*Aspergillus* IgG levels in sera. Consequently, there are fewer opportunities for immunodiffusion testing [14]. In Japan, the immunodiffusion method for detecting *Aspergillus* antibodies has been discontinued commercially, which has caused significant difficulties in diagnosing aspergillosis in penguins. As described in a report by Cabana et al., an *Aspergillus* antigen (galactomannan) detection kit was not useful for the diagnosis of aspergillosis in infected penguins [3]. Therefore, there is a need to develop serological tests that are useful for diagnosing aspergillosis in animals, including penguins. Commercially available human IgG anti-*Aspergillus* antibody detection kits detect human IgG using mouse anti-human IgG monoclonal antibodies, but they do not detect IgG or IgY antibodies in other species. Commercially available anti-chicken IgY antibodies show cross-reactivity with avian species including penguins [5]. In this study, we examined the combined use of anti-chicken IgY conjugated with horseradish peroxidase and an anti-*Aspergillus* IgG detection ELISA kit for the detection of anti-*Aspergillus* IgY in penguin sera.

## MATERIALS AND METHODS

### Ethical considerations regarding the use of animals

Health checks and all diagnostic and treatment procedures were conducted following the practice guidelines of the veterinary department of each aquarium. Experiments conducted using serum samples from penguins were approved by the Animal Care and Use Committee of Obihiro University of Agriculture and Veterinary Medicine (No. 21-40, 22-109, 23-28, 23-142).

### Sera collected from captive penguins

Sera of King penguins (*Aptenodytes patagonicus*), African penguins (*Spheniscus demersus*), and Gentoo penguins (*Pygoscelis papua*) were kindly shared by Noboribetsu Marine Park Nixe (Hokkaido, Japan) and Asahikawa, Asahiyama Zoological Park and Wildlife Conservation Center (Hokkaido, Japan). The sera that were used and information related to penguins are presented in Table 1 and Supplementary Table 1. When a King penguin (individual ID = 1) presented with coughing (day 0), a blood sample was collected (serum ID = 1a). Treatments with voriconazole (per os) from day 0 to day 3 and amphotericin B (nebulizing) from day 1 to day 4 were administered. On day 9, a second blood sample (serum ID = 1b) was collected. The patient was suspected to have aspergillosis, but no diagnosis was confirmed. An African penguin (individual ID = 8) was confirmed to have died of aspergillosis caused by *Aspergillus fumigatus*. The penguin presented inappetence and weakened responsiveness to external stimuli 19 days before death (set as day 0). Antibacterial drug treatment was initiated on day 1. Blood samples (serum ID 8a and 8b) were collected on days 4 and 5. A blood sample (serum ID = 8c) was collected on day 11. Itraconazole treatment (per os) was initiated on that day, in addition to antibacterial treatment. On day 19, the penguin died. The penguin was not only pathologically determined to have aspergillosis but also microbiologically determined because *A. fumigatus* was isolated from the lung.

**Table 1.**
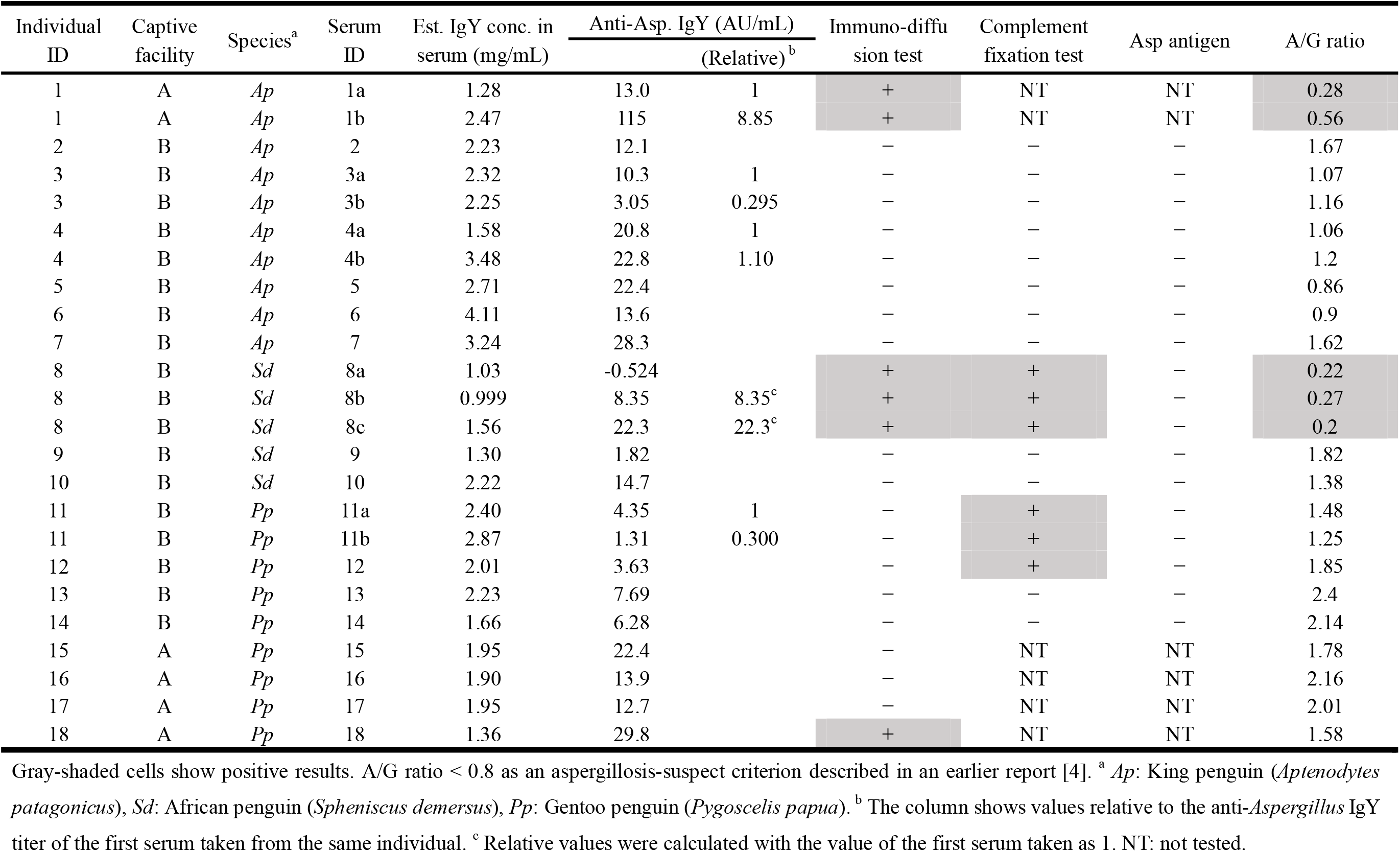
Overview of serologic tests related to the diagnosis of aspergillosis.

### Cross-reactivity between anti-chicken IgY antibody and penguin sera

To examine the cross-reactivity between anti-chicken IgY antibody and penguin sera, sheep anti-chicken IgY antibody conjugated with horseradish peroxidase (anti-chicken IgY-HRP) (Tokyo Chemical Industry Co., Ltd., Tokyo, Japan) was used in this study. Purified normal chicken IgY, which was used as a control, was purchased from Bio-Techne Corp. (Minneapolis, MN, USA). The procedure is as follows. MaxiSorp (Thermo Fisher Scientific, Waltham, MA) was used to coat the plates. Purified normal chicken IgY diluted with 50 mM bicarbonate/carbonate buffer (pH 9.6) was used as a serial standard and coated at 15.6, 31.3, 62.5, 250, 500, and 1000 pg in each well. Diluted penguin sera (100 µL) were put in each well After penguin sera were diluted 10^6^ fold with the bicarbonate/carbonate buffer because undiluted sera from penguins showed excessive reaction with 1000-fold diluted anti-chicken IgY (data not shown). After coating overnight at 4°C, the wells were washed three times with phosphate-buffered saline and then blocked with 1% BSA. After blocking at 25°C for 1 hr, the wells were washed thrice with 0.1% Tween-20 in Tris-buffered saline (TBS-T). Next, 100 μL of anti-chicken IgY-HRP diluted 1000 fold with TBS-T, following the manufacturer’s instructions, was added to each well. After incubation at 37°C for 1 hr, the wells were washed thrice with TBS-T. Next, 3,3′,5,5′-tetramethylbenzidine (TMB) substrate solution was added to each well. After incubation for 5 min at 25°C, 100 μL 1 M HCl was added to each well as a stop solution. Absorption at 450 nm and reference absorption at 620 nm were measured in each well.

### ELISA using Platelia *Aspergillus* IgG Kit and anti-chicken IgY antibody

Platelia *Aspergillus* IgG Kit (Bio-Rad Laboratories, Inc., Hercules, CA) and anti-chicken IgY-HRP (Tokyo Chemical Industry Co., Ltd.) were used for experimentation. Although almost all parts of the kit procedure were followed according to the manufacturer’s instructions, 1/1000 volume of anti-chicken IgY-HRP solution was added to the conjugate solution containing mouse anti-human IgG monoclonal antibody conjugated with peroxidase. After the kit stopping solution was added to each well, the absorbance was determined at 450 nm using a microplate reader (GENios Pro; Tecan Group Ltd., Männedorf, Switzerland).

### Serological immunodiffusion assay

*An Aspergillus* Immunodiffusion System (Microgen Bioproducts Ltd., Surrey, UK (now Gold Standard Diagnostics)) was used for the immunodiffusion (Ouchterlony) assay. For the assay, 0.4% Gelrite plates containing 0.1% sodium azide in McIlvaine’s citric acid phosphate buffer (pH 7.0) were prepared. After gel solidification, wells of 8-mm diameter were prepared. In each well, 40 μL of serum or positive antigen solution was used for the assay. The precipitin formation was confirmed every 24 hr for three days.

### Other serological tests

Electrophoresis, complement fixation test, and *Aspergillus* antigen (galactomannan) test were performed at an external testing facility (Daiichi Kishimoto Clinical Laboratories, Sapporo, Japan).

### Statistical analyses

Kruskal–Wallis tests followed by Scheffe’s method were used for multiple comparisons of the IgY concentrations. Mann–Whitney U-tests were used for comparison of the estimated titers of anti-*Aspergillus* IgY between the two facilities.

## RESULTS

### Penguin sera show cross-reactivity to anti-chicken IgY antibody

All sera from King, African, and Gentoo penguins used in this study showed higher absorbance at 450 nm than that of the control wells in the ELISA described in *Materials and Methods*. The IgY concentration estimated from a standard line of chicken IgY was 1.00–3.48 mg/mL (Table 1 and Figure 1). The estimated IgY concentrations in the sera used were significantly different between the King and African penguins (Figure 1). Although differences in the potency of cross-reactivity to anti-chicken IgY antibodies among the three penguin species are not known, the observed differences among penguin species might reflect the difference in cross-reactivity. These data demonstrate that the anti-chicken IgY antibody recognizes the components of the penguin serum. The main cross-reactive component is presumed to be IgY.

**Figure 1.**
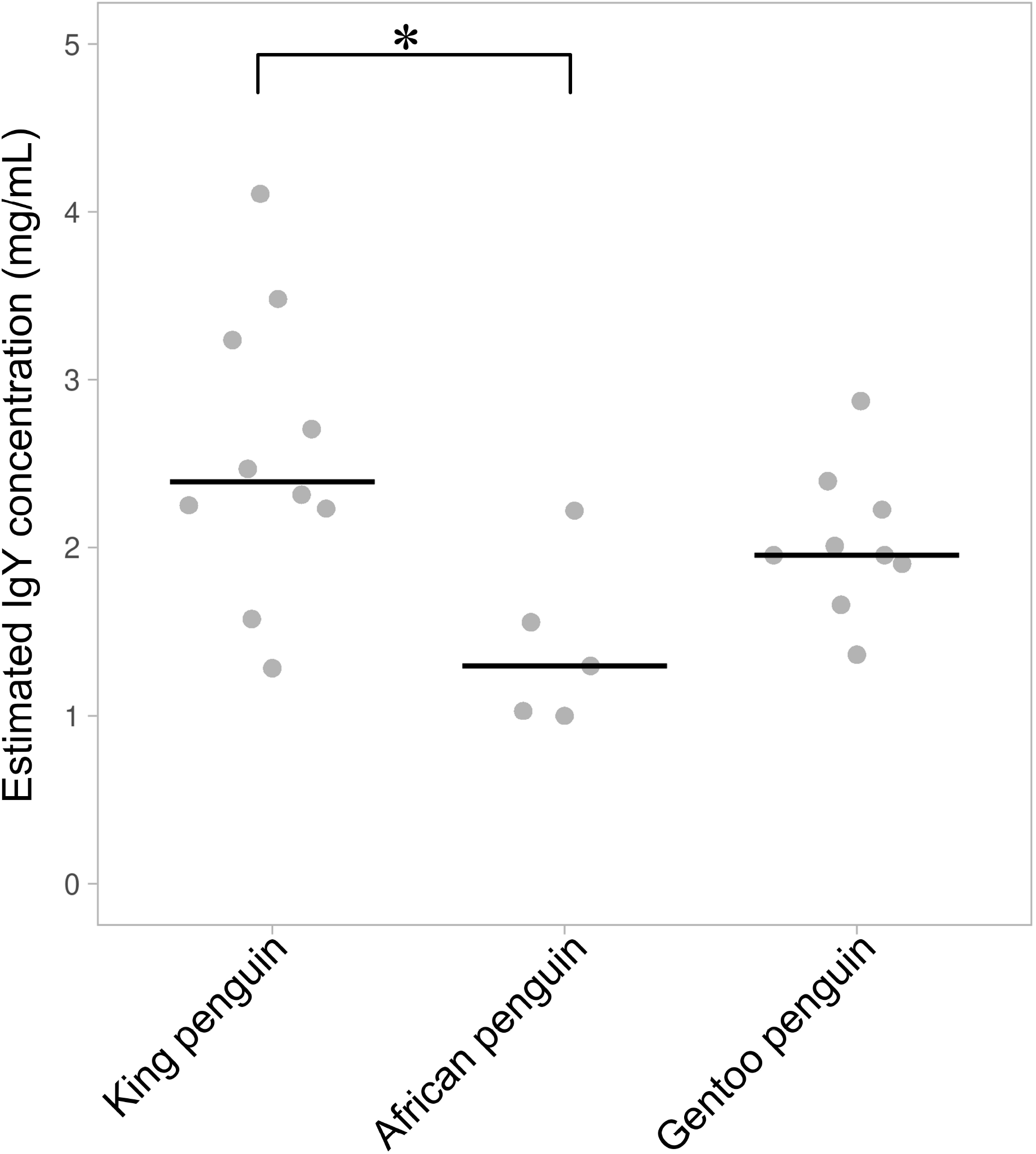
Estimated IgY concentrations in penguin serum samples. The plot was prepared using PlotsOfData (Postma and Goedhart 2019). Vertical lines represent the medians of each group. ∗ *p* < 0.05 using the Kruskal–Wallis test followed by Scheffe’s test as a post hoc test.

### Anti-*Aspergillus* titers in penguin sera are determined by anti-chicken IgY antibody

Serum samples were examined by ELISA to detect anti-*Aspergillus* IgY antibodies. The range of anti-*Aspergillus* titers was -0.5 to 115 eAU/mL (Table 1 and Figure 2 a). The titers increased over time in serial sera from individual ID 1 with suspected aspergillosis and eight with confirmed aspergillosis (Figure 2 a). The titers of two other serially collected sera samples (individual IDs 3 and 11) decreased over time. No major variation was observed in the serially collected sera from individual ID 4 (Figure 2 a). The titers of Gentoo penguins kept at facility B were higher than those of Gentoo penguins kept at facility A (Figure 2 b).

**Figure 2.**
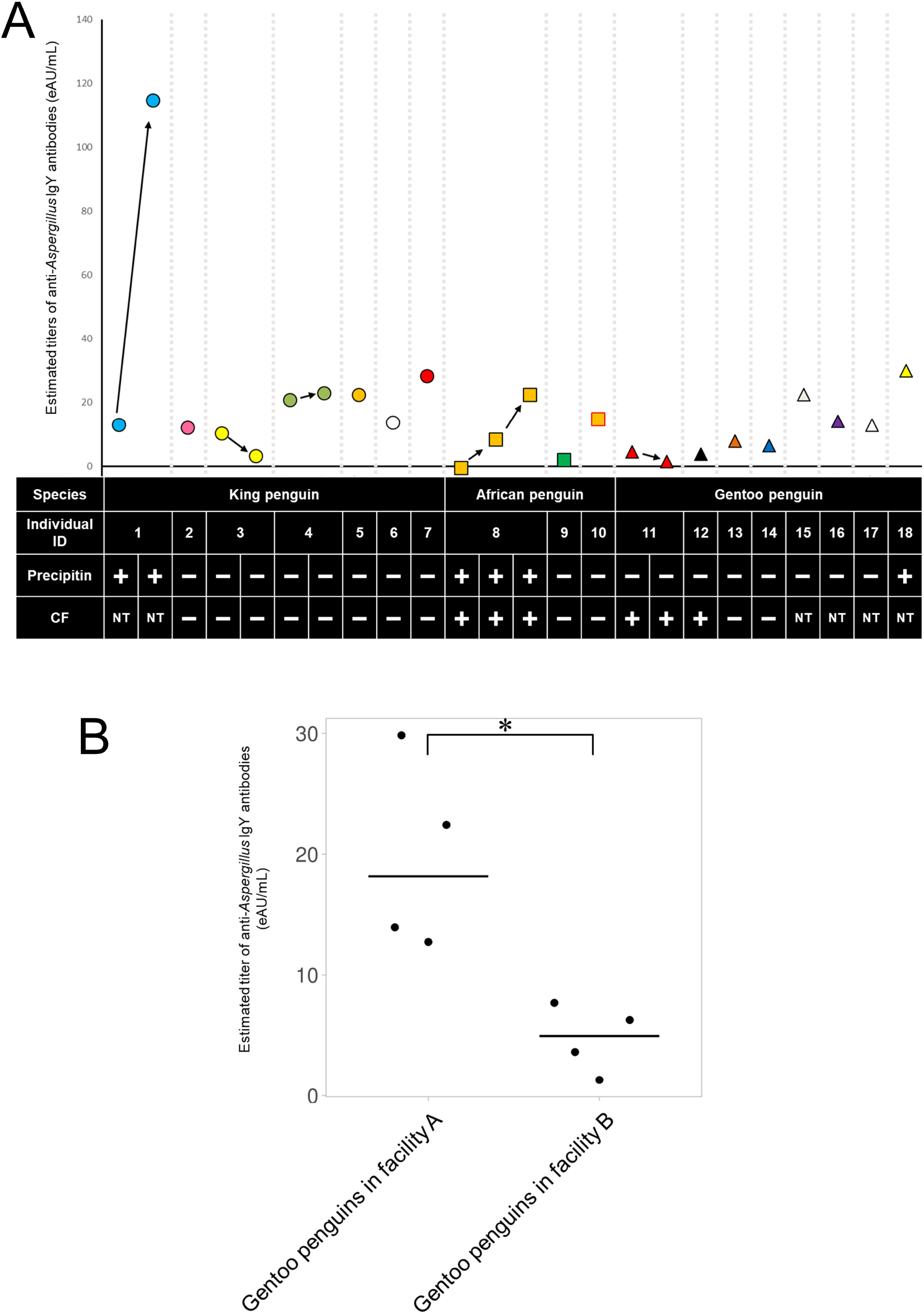
Estimated titers of anti-Aspergillus IgY antibody in penguin sera. (a) Overview of the titers of anti-*Aspergillus* IgY. Individual ID 1 was suspected of having aspergillosis. Individual ID 8 was confirmed as having aspergillosis. eAU means estimated AU: circles, King penguins; squares, African penguins; and triangles, Gentoo penguins. Results of the immunodiffusion assay and complement fixation are shown respectively in the rows of precipitin and complement fixation (CF). NT means not tested. (b) Comparison of the estimated titers of anti-*Aspergillus* IgY between two facilities. The plot was prepared using PlotsOfData (Postma and Goedhart 2019). Vertical lines represent the medians of each group. ∗ *p* < 0.05 using Mann–Whitney U-test.

### Results of precipitin formation, complement fixation, and Aspergillus antigen tests in serum samples

Precipitin formation was observed in the sera of an aspergillosis-suspected individual (ID 1) and an aspergillosis-confirmed individual (ID 8). Additionally, the serum from a healthy individual (ID 18) showed precipitin formation.

The complement fixation test was only performed at facility B. As shown in Table 1, three serum samples from two individuals (ID 11, 12) were found to be positive. However, the other indicators showed negative results.

*The Aspergillus* antigen (galactomannan) test was performed on the sera of penguins kept at facility B (Table 1). Even in serum samples from an aspergillosis case (individual ID 8), no *Aspergillus* antigen was detected.

## DISCUSSION

The diagnostic applications, including serological tests for aspergillosis in penguins, remain limited. In this study, we used an anti-*Aspergillus* IgG detection kit that combines an anti-chicken IgY antibody conjugated with HRP. Initially, we examined the cross-reactivity of anti-chicken IgY antibodies with penguin sera. In this study, we obtained serum samples from King, Gentoo, and African penguins. They showed higher absorbance than the control wells, indicating that they were reactive to the anti-chicken IgY antibody. These results were consistent with those described in previous reports [13, 15] As reported by Han et al., transcriptome analysis showed that Gentoo penguins retained only the IgY1 subclass and that chickens retained only the IgY2 subclass [9] It remains unknown why so much cross-reactivity is found in different subclasses, but the findings by Han et al. [9] that BLAST searches also indicated the possible presence of two IgYs in Gentoo penguins might help explain the cross-reactivity found with anti-chicken IgY antibodies. The concentrations were estimated as 1–3.5 mg/mL. The concentration of IgY in chicken sera is 3–7 mg/mL [17]. The concentrations were estimated based on the recognition of penguin IgY using anti-chicken IgY antibodies. The low measured values are likely due to the strength of the cross-reactivity. Next, we examined the titer of anti-*Aspergillus* IgY in penguin sera using a human anti-*Aspergillus* IgG measurement kit combined with an anti-chicken IgY antibody conjugated with HRP, as described above. In this study, serial serum samples of two penguins suspected (individual ID 1) or confirmed (individual ID 8) to have aspergillosis were included. Both cases showed that anti-*Aspergillus* IgY titers increased over time, indicating their potential for application in the diagnosis of aspergillosis. However, titers in healthy individuals were not constant. The cutoff values of the kit for human aspergillosis were 10 AU/mL and greater as positive and 5 to less than 10 as intermediate. Regarding the results obtained in this study, even healthy individuals showed positive results. Although the appropriateness of this cut-off value is not known because sufficient studies have not been conducted in penguins, these results suggest that the exposure levels of the penguins varied because *Aspergillus* spp. are common in the environment. It is particularly interesting that, among the Gentoo penguins, the titers of four individuals kept at facility A were higher than those of four individuals kept at facility B. A possible reason for this is related to differences in their respective environments. However, the reason for this finding remains unclear.

This study has several limitations. First, we employed a commercially available anti-chicken IgY antibody for serological testing. While its accessibility is advantageous, the cross-reactivity and specificity of this antibody toward penguin IgY may vary depending on several factors, including the penguin species, the quality of the polyclonal antibody, and lot-to-lot variability. These factors can influence the binding affinity and diagnostic performance, potentially affecting the accuracy and reproducibility of the results. Therefore, further validation is necessary to assess the antibody’s suitability for different penguin species and to establish reliable, species-specific cut-off values for penguin sera. Second, we observed considerable variation in background antibody titers, even among clinically healthy individuals. This heterogeneity complicates the interpretation of serological results and may lead to false-positive or ambiguous outcomes in the diagnosis of aspergillosis. In a recent study, an Aspergillus-specific IgG antibody test, which is not an ELISA test, showed a false-positive result in one-third of healthy human patients [18]. Therefore, continuous monitoring and refinement of diagnostic criteria are essential to enhance the reliability of early detection and improve treatment outcomes.

Among the serological examinations, the A/G ratio and precipitin formation in immunodiffusion tests were the most reliable indicators of aspergillosis in this study. In the future, immunodiffusion tests may become less commercially available. Therefore, measurement of the A/G ratio is important for recognizing aspergillosis. However, the A/G ratio is not specific to aspergillosis. Therefore, respiratory abnormalities must be determined by CT imaging when available. Furthermore, to increase specificity, determination of anti-*Aspergillus* IgY titers may be useful. As shown in the examination, the background titer level depends on the surrounding environment. Determining the usual titer of each might be beneficial to detect the change in the titer of anti-*Aspergillus* IgY for prompt treatment against aspergillosis.

## ACKNOWLEDGMENTS

This research was supported by the Japan Society for the Promotion of Science KAKENHI (C), Grant Number 21K05931. We thank the staff of Noboribetsu Marine Park Nixe and Asahikawa, Asahiyama Zoological Park and Wildlife Conservation Center for their cooperation with this study.

